# A Cell-Type–Resolved Meta-Analysis Reveals Glial DNA Methylation Changes Associated with Aging and Alzheimer’s Disease

**DOI:** 10.64898/2026.05.04.722662

**Authors:** Uchit Bhaskar, Mark Z Kos, Melanie A. Carless

## Abstract

Epigenome-wide association studies implicate DNA methylation in the development and progression of Alzheimer’s disease (AD). Although recent studies show that the epigenetics of non-neuronal cell types contribute to disease risk, the role of the methylome in individual glial cell types (i.e., astrocytes, oligodendrocytes) in biological aging and AD pathogenesis is unclear. In this study, we examined archived DNA methylation data across 13 cohorts and performed cell type deconvolution *in silico* to identify novel epigenetic signatures associated with aging and AD in glial cells. We observed pronounced age-associated methylation in astrocytes within the prefrontal cortex, whereas oligodendrocytes of the entorhinal cortex show the most differential methylation with AD status. Astrocytes, along with neurons, within the prefrontal cortex, emerge as key players in Braak stage-associated methylation, exhibiting strong concordance with previously reported associations at the brain tissue level. Age-associated changes in oligodendrocytes exhibit strong directional correlation with, and amplification of age-related effects with AD that affect neurodevelopmental processes, while AD-related methylation changes at age-associated sites in astrocytes diverge from those representative of normative aging processes. Our study expands on previous findings and reveals glial-specific methylation patterns associated with epigenetic aging and AD.

## Introduction

Alzheimer’s disease (**AD**) is largely characterized by the pathological accumulation of amyloid and tau proteins, leading to neuronal loss and dysfunction, which manifests as progressive cognitive decline. Recently, attempts have been made to unravel the role of glial cell types (including astrocytes and oligodendrocytes) in disease progression, including dysfunctions in maintaining CNS homeostasis, blood-brain barrier integrity, myelination and axonal support, metabolism, and synapse maintenance.^1–4^ Several epigenome-wide association studies (**EWAS**) have pointed to a role for DNA methylation in the etiology of AD and show differential DNA methylation across several CpG sites in relation to pathogenesis.^5–19^ Recently, studies have shown that among differentially methylated positions (**DMPs**) associated with Braak stages, a large fraction seems to stem from non-neuronal (NeuN^-^) cell types,^15^ and underscored differences in neuronal and non-neuronal CpG methylation in relation to AD,^7^ highlighting a critical need to resolve cell-specific epigenetic contributions to disease states.

To date, most studies profiling DNA methylation in AD utilize bulk brain tissues, rather than single-cell profiling, especially for glial subtypes. An alternative approach to assess cell-type-specific epigenetic contributions is to use computational deconvolution methods.^20–23^ One recent study used tensor-composition-based deconvolution analysis and identified a novel promoter hypomethylation signature at the *PEN-2* gene, associated with amyloid plaque burden in neurons.^24^ However, further exploration is necessary to define changes in the glial epigenome in relation to both age and disease pathology.

In this study, we examined archived DNA methylation data from 13 different cohorts, totaling ∼3,812 samples and spanning several different brain regions, to identify cell-type-specific methylomic variation associated with age and AD. We show that a significant proportion of DMPs associated with age occur in glial cell types (i.e., astrocytes and oligodendrocytes), as well as endothelial cells. AD associations tend to be highly specific to cell type and brain region. We observed a significant proportion of DMPs associated with AD status in oligodendrocytes compared to other cell types, particularly in regions affected early in the disease state (e.g., entorhinal cortex). DMPs within the PFC, previously identified to be associated with Braak stage, are preferentially enriched in both astrocytes and neurons. Importantly, age-associated DNA methylation signatures in oligodendrocytes across all cortical regions tend to be exacerbated with AD pathology, whereas age-associated signatures of astrocytes seem to be decoupled with disease state. Together, this suggests the existence of distinct cell-specific epigenetic programs in both healthy aging and neurodegeneration.

## Results

We analyzed cell-type-specific DNA methylation differences associated with age and AD-status across a total of 7 brain regions – prefrontal cortex (**PFC**), middle temporal gyrus (**MTG**), superior temporal gyrus (**STG**), entorhinal cortex (**EC**), the broad temporal cortex (**TC**), hippocampus (**HC**), and the cerebellum (**CBL**) (**Fig. 1a**). Since prior EWAS pointed to roles for non-neuronal cell types in disease pathology, and age is the largest risk factor for AD, we were interested in evaluating regional glial cell vulnerability associated with both aging and AD. For our initial analysis, we focused on either age as a continuous variable or AD status as a binary variable, since stage-specific disease data were not available across all cohorts. However, for additional analysis, we also used Braak stage as a continuous variable to assess cell-specific associations when available. We employed a cell-type decomposition method, using an existing *EpiSCORE* brain reference panel,^25,26^ for each study and performed cohort-wise, region-stratified, cell-specific association analyses using CellDMC for age, AD, and Braak stages.^27^

**Figure 1.**
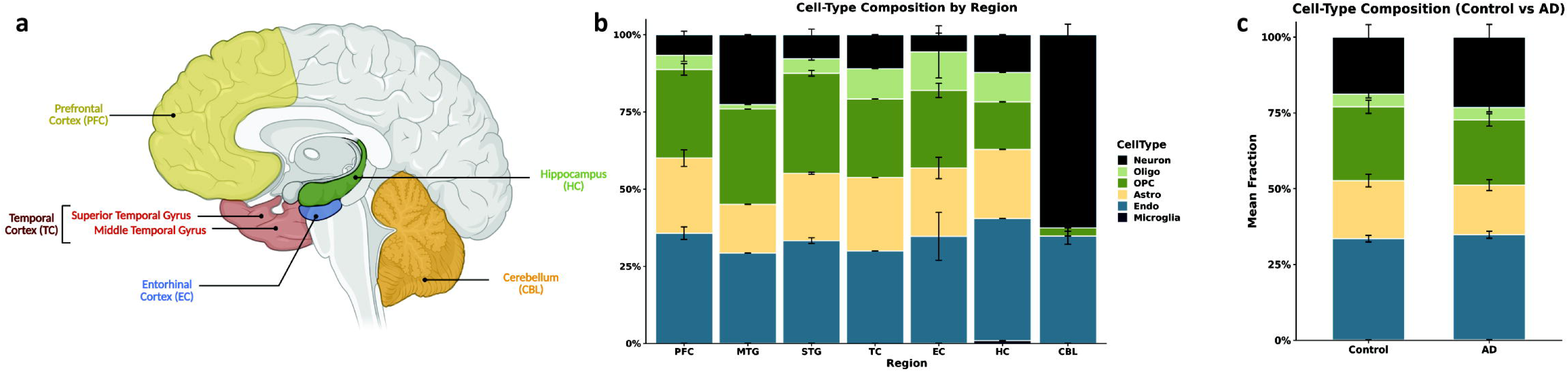
Overview of brain regions and cell types used in the study. **a)** Representative schematic of the human brain, highlighting brain regions analyzed in this study. Cell-type composition estimates by **b)** region and **c)** disease status. The sizes of stacked bars represent mean fractions across all cohorts analyzed. Error bars represent standard errors.

Cell type compositions were largely similar across multiple cortical regions (**Fig. 1b**), irrespective of healthy and disease states (**Fig. 1c**). Overall, across cortical and hippocampal tissues, neurons accounted for 9.53±1.36% of total cells, astrocytes 21.41±0.94%, endothelial cells 36.34±1.66%, oligodendrocytes 8.03±1.82%, oligodendrocyte precursor cells (OPCs) 24.33±2.09%, and microglia < 0.6% (0.35±0.17%). In the CBL, we saw a distinct pattern, with only neurons (60.4±2.53 %), oligodendrocytes/OPCs (3.24±0.79 %), and endothelial cells (36.19±1.95%) making up the cell types. For each cohort, since oligodendrocytes represented only a small fraction, we grouped these with OPCs, making up a common “Oligo_OPC” pool (hereafter, referred to as oligodendrocytes for simplicity), and removed cell types existing in less than 0.5% of the samples (i.e., microglia in most cohorts). Overall, we leveraged power by performing a fixed-effect inverse variance weighting (**IVW**)-based meta-analysis across 13 cohorts (n = 3,801 unique samples, *see Methods*) to uncover novel cell-type-specific differentially methylated loci for age and AD.

### Cell-Type Deconvolution Uncovers Unique Glia-Specific Age-Associated DMPs

Age is a significant risk factor for the development of several neurodegenerative disorders. Previously, epigenetic age acceleration has been shown to be associated with AD-related pathology, including neuritic plaques and neurofibrillary tangles,^28^ suggesting that age-related methylation might be a key player in mediating neurodegenerative phenotypes. Importantly, identifying age-associated methylation in a cell-type-specific manner will help us better understand cell-type vulnerability patterns associated with late-stage brain disorders. Since cell-type deconvolution methods substantially reduce statistical power due to the high dimensionality of the data, we chose to correct for multiple testing and assess significance using a false discovery rate (FDR; Benjamini-Hochberg) threshold of 0.05. With this, we identified 2,101 significant DMPs, of which neurons, astrocytes, oligo-OPCs, and endothelial cells represented 37.93 %, 32.27 %, 11.09% and 18.71% of total hits, respectively (**Fig. 2a**). Neurons, accounting for the largest population in the cerebellum, also contributed to most (∼63%) of the significant age-associated cerebellar DMPs; whereas glial cells dominated cortical tissues (**Fig. 2b**).

**Figure 2.**
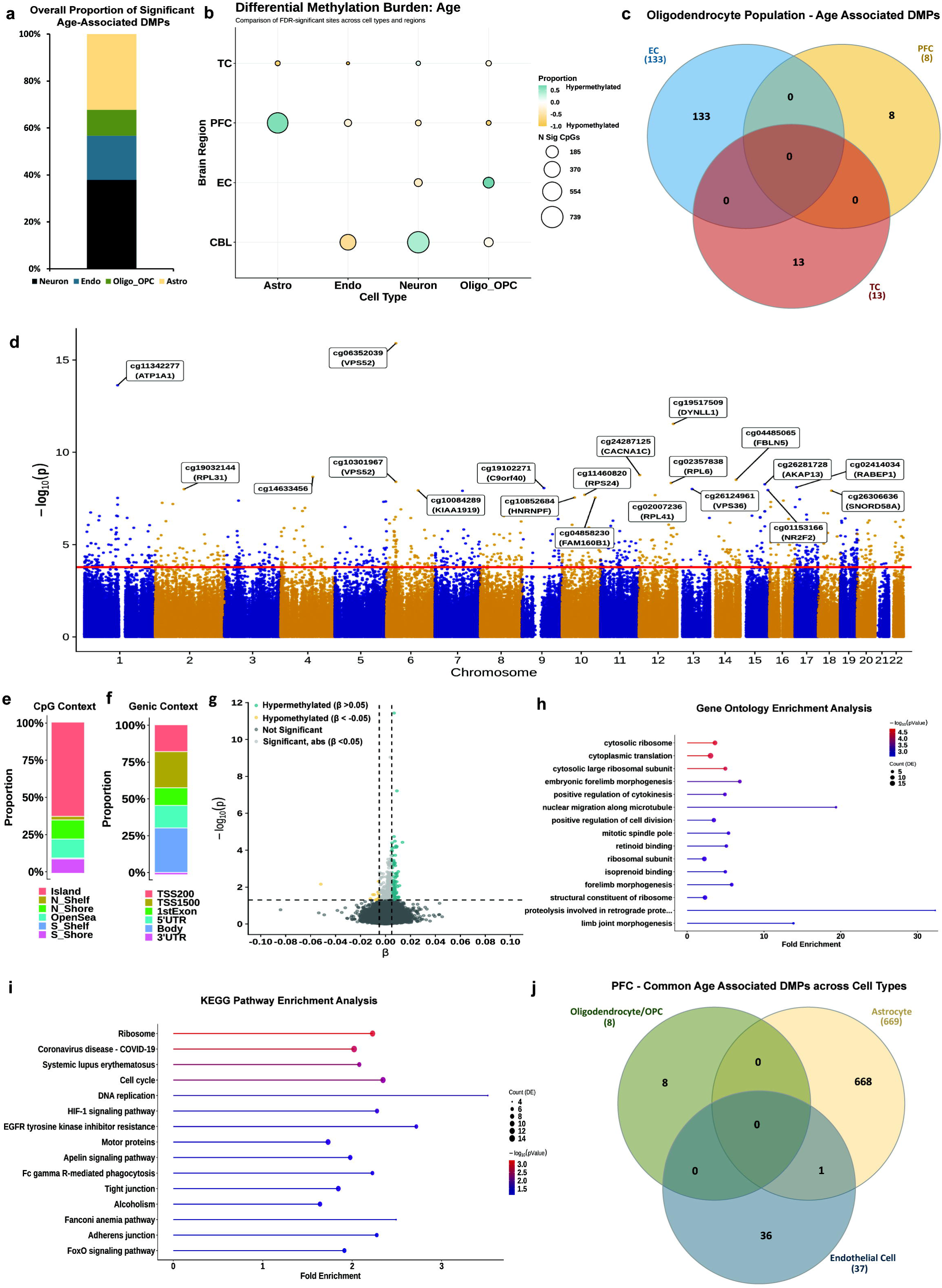
Region-stratified cell-type deconvolution-based meta-analysis identifies glia-specific aging programs. **a)** Stacked bar chart showing overall proportions of significant age-associated differentially methylated positions (DMPs) across all regions. Significance set at FDR < 0.05. **b)** Bubble plot showing differential methylation burden associated with age across all cell types x regions analyzed. Color scale represents proportion of sites seen to be hypermethylated (teal) and hypomethylated (orange). Size of bubbles indicates number of significant CpGs. Also see Supplementary Fig. S1. **c)** Venn diagram of age-associated DMPs in the oligodendrocyte fractions across all cortical regions, showing no shared loci. **d)** Manhattan plot for the meta-analysis on prefrontal cortex astrocytes (N = 1,626 individuals), with the 20 most significant DMPs annotated with either CpG ID or Illumina UCSC gene name or both. X-axis shows chromosomes 1-22 and y-axis shows -log_10_(p), with the horizontal red line denoting FDR significance (0.05). **e)** Stacked bar chart showing the proportion of significant age-associated DMPs in PFC astrocytes that are enriched for different CpG contexts, based on Illumina UCSC annotations. **f)** Stacked bar chart showing the proportion of significant age-associated DMPs in PFC astrocytes that are enriched for different genic positions, based on Illumina UCSC annotations. **g)** Effect size volcano plot of age-associated DMPs in PFC astrocytes. Horizontal line identifies FDR threshold of 0.05, whereas the vertical lines indicate effect size, |β| > 0.05. **h)** Lollipop plot of the 15 most enriched GO terms for age-associated DMPs in PFC astrocytes, ordered by counts. X-axis represents fold-enrichment and color scale represents -log_10_(p). **i)** Lollipop plot of 15 most enriched KEGG pathway analysis terms for age-associated DMPs in PFC astrocytes, ordered by highest counts. X-axis represents fold-enrichment and color scale represents -log_10_(p). **j)** Venn diagram of common age-associated DMPs across astrocytes, oligodendrocytes, and endothelial cells within the PFC.

Astrocytes harbored 93% of the significant DMPs in the PFC (669 CpG sites) and 9 DMPs in the TC. Oligodendrocytes held 71.5% of the significant DMPs in the EC (133 CpG sites), 1.1% in the PFC (8 DMPs), and 4.64% in the TC (13 DMPs). We did not observe any intersection of the oligodendrocyte-significant DMPs across the three cortical regions (**Fig. 2C**), indicating brain region-specific CpG methylation patterns with age among these cells. Of the 133 DMPs in the EC oligodendrocytes (**Supplementary Fig. S2a**), 113 are hypermethylated with age, and 20 sites are hypomethylated. Among the top 10 hits, we observed hypermethylation at a site within the *DCDC2* gene body (cg11563656, ES = 0.016764436, p = 3.93 x 10^-10^), previously identified as a predictor for episodic memory maintenance in aged adults.^29^ Gene Ontology (GO) analysis showed enrichment for neurodevelopmental processes, including Notch signaling, and those involved in protein metabolism, to be associated with age in these cells (**Supplementary Fig. S2b**).

Since the PFC represented our largest sample set, and astrocytes showed the highest number of significant DMPs in this region, we focused on these cells. Of the 669 DMPs, 567 sites mapped to known genes (∼84.7%) in the genome, including novel associations at three sites within the *VPS52* gene (cg06352039, p = 1.24 x 10^-16^; cg09112371, p = 6.75 x 10^-6^; cg10301967, p = 3.94 X 10^-9^), as well as hits at genes with known astrocyte functions such as ionic and glutamate transport (*ATP1A1*, cg10442913, p = 2.95 x 10^-8^; cg13079047, p = 2.87 x 10^-7^), calcium signaling (*CACNA1C*, cg24287125, p = 1.73 x 10^-9^), and astrocyte reactivity (*FBLN5,* cg04485065, p = 3.06 x 10^-9^) (**Fig. 2d**). We also observed other significant DMPs mapped to genes of the SLC, ADAM and KLF families (**Supplementary Table S1**), and a site mapped to the *PRDM2* gene (cg23813012, p = 8.54 x 10^-7^) that was previously observed in an age-association EWAS of sorted neuronal and non-neuronal cell types.^7^

Of the 669 identified significant DMPs, nearly 60% are located within CpG islands, about 25% in adjacent CpG “shore” regions (within 2 kb of islands), and the remainder in CpG “shelves” (2-4 kb) and “open sea” regions (> 4kb) (**Fig. 2e**). Over a fourth of the sites map to known gene bodies, and nearly 40% of sites map to regions upstream of the transcription start site (most between 200-1,500 base pairs upstream) (**Fig. 2f**). A smaller fraction of the sites map to either 5 UTRs (∼15%) or first exons (∼9%). Of the 669 significant DMPs, most showed small effect sizes (ES), with only 13 hypomethylated sites having an ES < -0.005 and 114 hypermethylated sites having an ES > 0.005 (**Fig. 2g)**. The highest ES was observed for cg14633456 (ES = 0.0136, p = 2.16 x 10^-9^), a site previously implicated in aging.^30,31^ Gene ontology (**Fig. 2h**) and KEGG pathway analysis (**Fig. 2i**) showed significant enrichment (p < 0.05) for GO terms related to translation and protein processing (e.g., cytosolic ribosome, cytoplasmic translation), cell division (e.g., positive regulation of cytokinesis and cell division, cell cycle, DNA replication), as well as those involved in phagocytic and inflammatory processes, all of which are known to be disrupted with age. Differentially methylated region (**DMR**) analysis in the PFC-astrocytes identified 224 significant regions (Sidak corrected p < 0.05), with the most significant DMR mapping to the *ATP1A1* gene (Sidak p = 1.66 x 10^-21^) (**Supplementary Table S2**). Sixty-five of the significant PFC-astrocyte loci are also nominally significant in the TC (**Supplementary Fig. S3a**), with a significantly positive ES correlation (r = 0.78, p = 8.77 x 10^-15^), suggesting potential shared mechanisms for age-related changes across the cortex in astrocytes (**Supplementary Fig. S3b**). Finally, we assessed whether any of the sites significantly associated with age in astrocytes were also implicated in other cell types in the PFC (**Fig. 2j**). In the oligodendrocyte pool, only 8 sites showed significant associations with age (FDR < 0.05), none of which were common to those seen in astrocytes; and in endothelial cells only 1 of the 37 age-associated sites was observed among the astrocyte DMPs. We did not detect any significant association of PFC neurons with age.

### Oligodendrocytes exhibit region-specific differential methylation associated with Lipid Metabolism in AD

We performed association analysis using AD status as a binary variable, while controlling for age and sex in a region-stratified manner. Like our age-association study, we utilized a fixed-effect IVW method to meta-analyze these effects across the PFC, TC, EC, and CBL. Overall, in comparison to our age meta-EWAS, we observe far fewer significant associations with AD status across all regions and cell types (20 in total) at a Bonferroni significance threshold of 0.05 (p = 1.25 x 10^-7^ for ∼400,000 sites) (**Supplementary Fig. S4a**). With an FDR threshold of 0.05, we observe only 36 significant hits across all cell types in the PFC, 20 hits in the CBL, and only one significant DMP in the TC (**Fig. 3a**). However, AD-association in the EC shows a significantly higher proportion of DMPs (283 significant sites across all cell types, **Supplementary Fig. S4b**), indicating region-specific differences in DNA methylation associated with AD. As with age, glial cells, and to some extent, endothelial cells dominated DMPs in the cortical regions.

**Figure 3.**
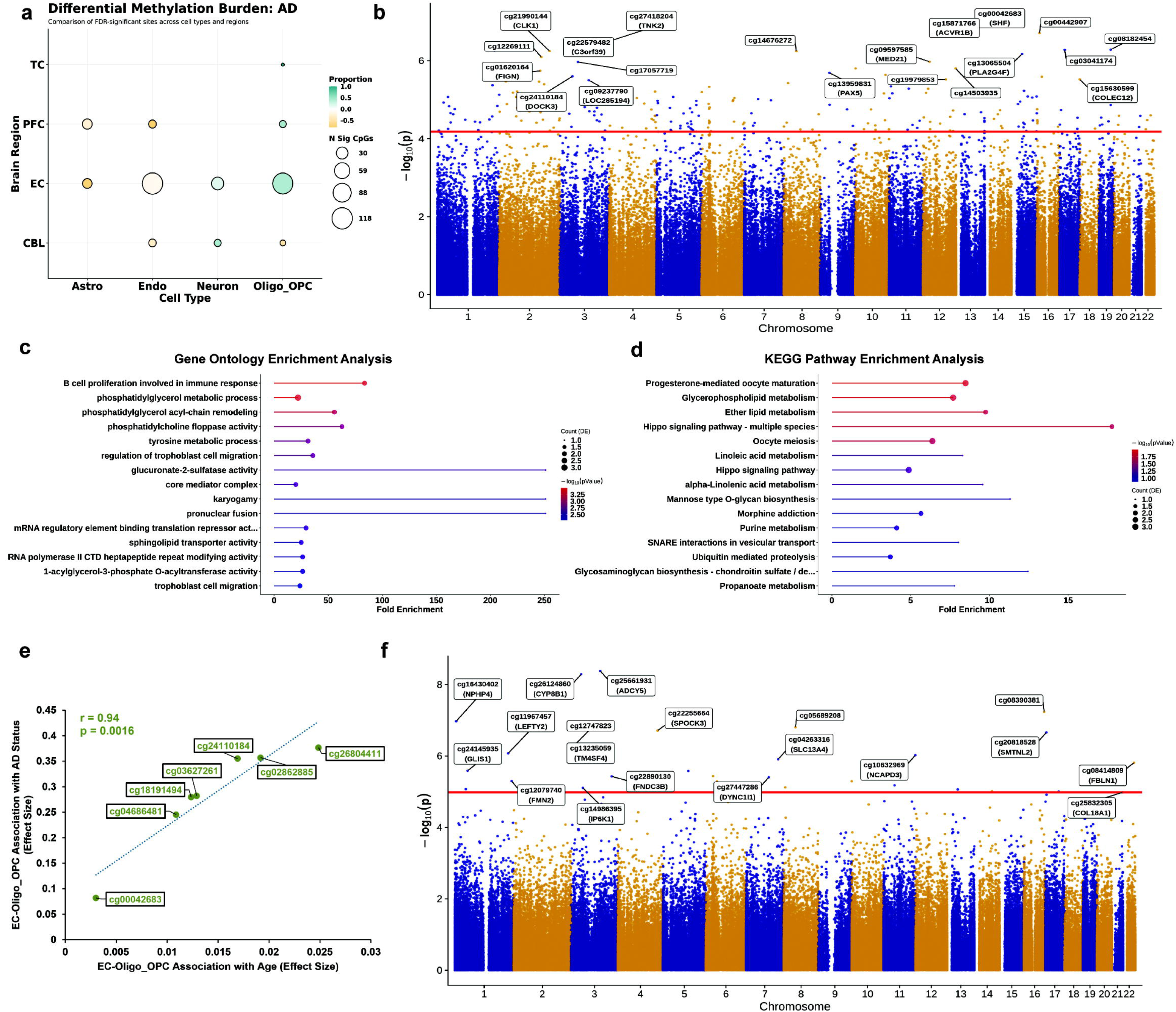
AD-associated DMPs exhibit region- and cell type specificity. **a)** Bubble plot showing differential methylation burden associated with AD status across all cell types x cortical regions analyzed. Color scale represents the proportion of sites seen to be hypermethylated (teal) and hypomethylated (orange). The size of bubbles indicates the number of significant CpGs. **b)** Manhattan plot for the meta-analysis on entorhinal cortex oligodendrocytes (N = 154 individuals), with the 20 most significant DMPs annotated with either CpG ID or Illumina UCSC gene name or both. X-axis shows chromosomes 1-22 and Y-axis shows -log_10_(p), with the horizontal red line denoting FDR significance (0.05). Also see Supplementary Fig. S5. **c)** Lollipop plot of the 15 most enriched GO terms for AD-associated DMPs in the EC oligodendrocytes, ordered by counts. X-axis represents fold-enrichment and color scale represents -log_10_(p). **d)** Lollipop plot of the 15 most enriched KEGG pathway analysis terms for AD-associated DMPs in EC oligodendrocytes, ordered by counts. X-axis represents fold enrichment. Color scale represents -log_10_(p). **e)** Correlation scatter plot of significant DMPs shared between age- and AD-associations in EC oligodendrocytes (r = 0.94, p = 0.0016, assessed by t-test). **f)** Manhattan plot for the meta-analysis on prefrontal cortex astrocytes (N = 1,626 individuals), with the 19 most significant DMPs annotated with either CpG ID or Illumina UCSC gene name or both. X-axis shows chromosomes 1-22, and y-axis shows - log_10_(p), with the horizontal red line denoting FDR significance (0.05).

EC oligodendrocytes exhibit a substantial proportion of differentially methylated sites associated with AD. Of these, 80 sites are hypermethylated with AD, while 34 sites are hypomethylated (**Fig. 3b and Supplementary Fig. S4b**). Our most significant hit is for a CpG site located within the gene body of the *SHF* gene (cg00042683, ES = 0.082, p = 7.65 x 10^-8^), known to play a critical role in oligodendrocyte fate specification and found to be downregulated in AD.^32^ Critically, we see hypermethylation at a novel CpG site in the 3’ UTR of the *ABCA1* gene (cg13559409, ES=0.24, p = 1.77x10^-5^), a key player of cholesterol transport, and lipidation of the well-known late-onset genetic risk factor, *APOE*.^33–36^ Other notable significant AD-related DMPs include *DOCK3* (a presenilin binding protein), *PDE4D* (calcium modulator impacting memory), and *SERPINI1* (important for limiting excitotoxicity) (**Supplementary Table S3**). Gene Ontology and KEGG pathway enrichment analysis for DMPs in the EC oligodendrocytes identifies several terms relevant to fatty acid and lipid metabolism-related processes, as well as those involved in proteoglycan biosynthesis (**Fig. 3c-d**), supporting a role for DNA methylation in modulating dysregulation of functional oligodendroglial cell states. Of note, DMR analysis of these results shows significant hits at a neuroinflammatory gene, *C1QTNF7* (Sidak corrected p = 4.59 x 10^-7^), containing 7 sites, as well as 3 sites within the *PLEC* gene (Sidak corrected p = 7.03 x 10^-5^), previously implicated as a critical player in pathological tau accumulation (**Supplementary Table S4**).^37,38^ Our study now strengthens these findings by implicating a role for abnormal DNA methylation in these processes.

Since we observed several significant hits associated with age in EC-oligodendrocytes, we asked whether any of these sites are shared between age and AD, and if so, how they are correlated. A total of 7 CpG sites are significantly associated with both age and AD in EC oligodendrocytes (**Fig. 3e**). While all seven DMPs are hypermethylated with both age and AD with a Pearson’s correlation of 0.94 (p = 0.0016, Student’s t-test), effect sizes for individual sites are much higher in AD associations (in the range of 0.08 – 0.38), than in the age-association (range of 0.003 – 0.02). These loci occupied multiple genes, which have previously been implicated in AD-related pathogenesis, and include our most significant AD hit within the *SHF* gene body (cg00042683), as well as sites within gene bodies of *DOCK3* and *PDE11A* (**Supplementary Table S5**). The substantial increase in effect size with AD at these sites points to disease-specific exacerbation of age-related methylation changes, likely specific to oligodendrocytes of the EC. Interestingly, to our knowledge, no previous EWAS of AD has reported a significant association for these sites, therefore, highlighting plausible novel candidates of AD-related age exacerbation in oligodendrocytes. We did not observe any shared significant sites in the oligodendrocyte population from the other cortical regions (PFC and TC).

Within the PFC, similar to the age-associated methylation patterns, astrocytes showed the highest number of DMPs. Of 36 significant DMPs across all cell types, 19 sites are detected in astrocytes (**Fig. 3f**), of which 7 are hypermethylated, and the remaining 12 are hypomethylated (**Supplementary Table S6**). Our most significant DMP is at a site (cg25661931) within a CpG island in the first exon of the *ADCY5* gene (ES = -0.091, p = 4.16 x 10^-9^). *ADCY5* encodes one of the adenylate cyclase family of cAMP producers and has previously been linked to the accumulation of hyperphosphorylated tau in AD.^39^ Also, cg25661931 within *ADCY5* has been linked to an epigenetic switch point associated with aging in a recent study.^40^ Differential methylation at the *HOXA3* gene cluster has been associated with AD pathology in the PFC.^7,17,18^ In astrocytes, we identified a nominally significant DMP (ES=0.12, p=0.008) within this gene (cg19048532). Nevertheless, most of the DMPs identified in the PFC-astrocytes showed only low effect sizes in association with AD, and only two sites were shared with age-associated DMPs in PFC-astrocytes. We also did not observe any significant shared DMPs between astrocytes in the EC and PFC with disease status (data not shown). However, DMR analysis identifies 20 significant regions associated with AD in PFC astrocytes (**Supplementary Table S7**). Of these, a significantly associated DMR is present within *SLC5A5* (Sidak corrected p = 0.001, 7 CpG sites), an ion transport gene previously shown to be protective for AD and frontotemporal dementia.^41,42^

### Astrocytes and neurons drive methylation signals associated with Braak stages

Although Braak stage data were not publicly available for all cohorts in our analysis, we conducted a meta-analysis for cohorts that had this data available. In total, we were able to assess Braak stage associations in the PFC for 1506 samples (across 6 cohorts), and in the TC (combined STG, MTG, and broad TC) for 296 samples (3 cohorts) (**Supplementary Table S8**). Of the three cohorts that had EC data available, only two provided Braak stage as a phenotype, and one of these did not meet model rank requirements, so we excluded the EC from our analysis. We observed a total of 10 significant DMPs in the PFC, and 9 in the TC, at an FDR adjusted threshold of 0.05 (**Fig. 4a**). Specifically, within the PFC, we observed 3 DMPs in astrocytes, 6 in the oligodendrocytes and 1 in neurons, whereas, in the TC, we saw 2 DMPs each in the astrocytes and oligodendrocytes, 1 in endothelial cells and 4 in the neurons. There was no overlap of significant DMPs between AD (as a binary trait) and Braak staging association analyses (data not shown). DMR analysis indicated cell-type-specific methylation signatures in the PFC, implicating 32 significant (Sidak p < 0.05) astrocyte-associated DMRs, 16 oligodendrocyte-associated DMRs, 13 neuron-associated DMRs, and 2 endothelial cell-associated DMRs (**Supplementary Table S9**). Our most significant DMR in PFC astrocytes is located in the 3 UTR of the *DLX1* gene (Sidak p = 2.75 x 10^-5^), previously associated with neuronal tau accumulation in progressive supranuclear palsy (**PSP**).^43^ Notably, we also observed an astrocyte-associated DMR on the *HOXC4* gene, containing 5 probes (Sidak p = 0.024), uncovering potential additional roles for the HOX clusters in AD pathology. DMR analysis of the temporal cortex showed 17, 19, 16, and 19 significant DMRs associated with Braak stage in astrocytes, oligodendrocytes, neurons, and endothelial cells, respectively (**Supplementary Table S10**). In the oligodendrocyte pool, we noted a DMR on a different HOX gene, *HOXC8* (Sidak p = 2.52 x 10^-5^), as well as multiple DMRs on a zinc-finger protein, *ZIC4*. We did not observe any significant cell-type-specific DMRs within some previously identified popular Braak stage-associated genes, including those within the HOXA clusters and ANK1, but identified one on the *CDH1* gene in astrocytes (Sidak p = 0.014) and another on the *KCNQ1* in neurons (Sidak p = 0.001), genes which were previously identified to house DMPs in association with Braak stage.^7^

**Figure 4.**
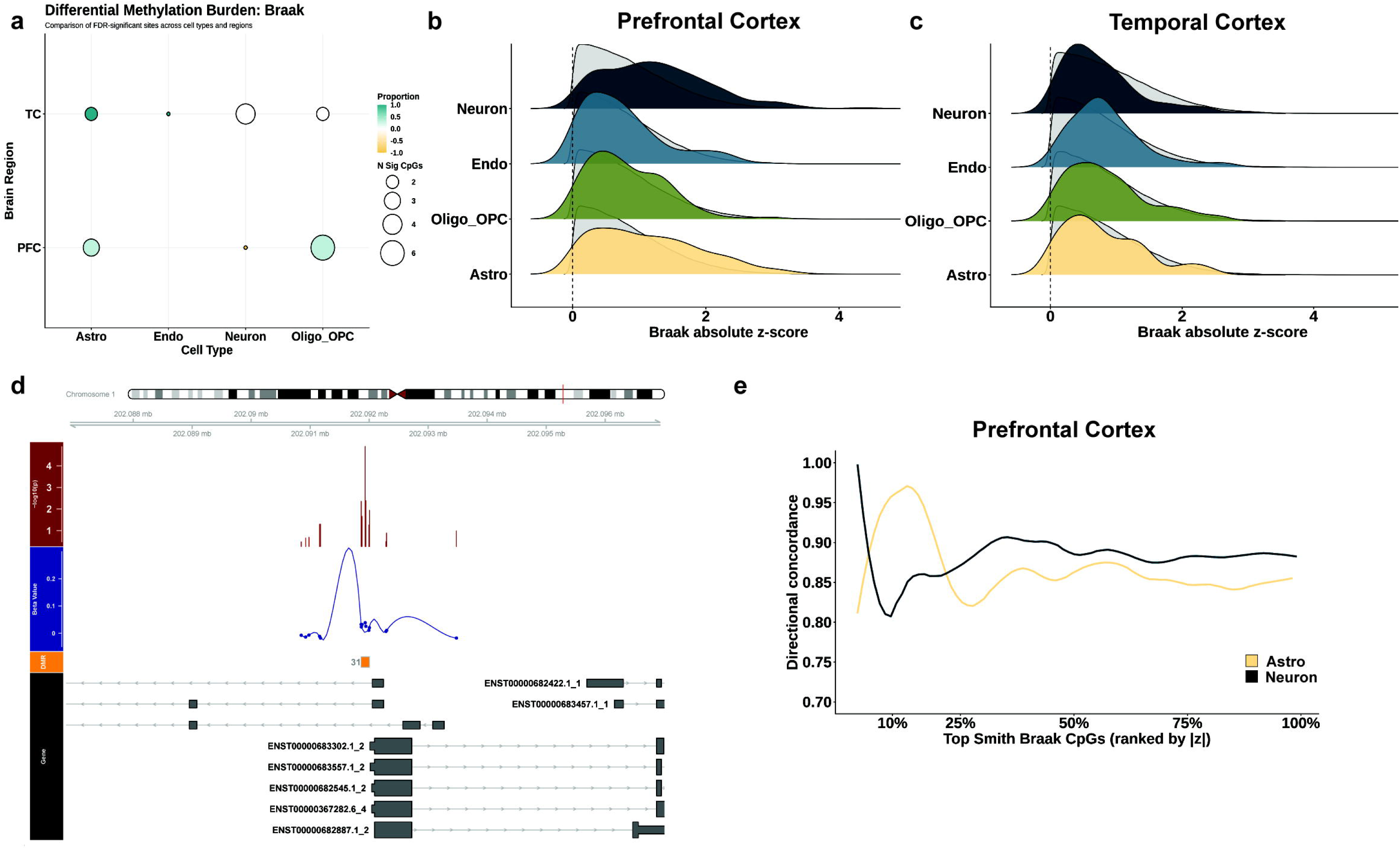
Astrocytes and neurons show distinct differential methylation trajectories in association with Braak stages. **a)** Bubble plot showing differential methylation burden associated with Braak stage across all cell types x cortical regions analyzed. Color scale represents proportion of sites seen to be hypermethylated (teal) and hypomethylated (orange). Size of bubbles indicates number of significant CpGs. **b)** Representative ridge plots for rank-based enrichment analysis of effect sizes for bulk tissue Braak-associated CpGs, identified in Smith et. al.,^18^ across all cell types in the prefrontal cortex. The X-axis shows absolute z-scores, defined as ES / SE for Braak associations of each cell type in the meta-analysis. For each cell-type, absolute z-scores of all ∼400k CpG sites are ranked, and probability distributions are shown as grey shaded ridges. 236 Braak-associated CpGs identified as significantly associated in bulk PFC by Smith et al. are treated as a predefined reference set, and absolute z-score distributions of these sites are represented by colored ridges. **c)** Representative ridge plots for rank-based enrichment analysis of effect sizes for bulk tissue Braak-associated CpGs, identified in Smith et. al.,^18^ across all cell types in the temporal cortex. The X-axis shows absolute z-scores, defined as ES / SE for Braak associations of each cell type in the meta-analysis. For each cell-type, absolute z-scores of all ∼400k CpG sites are ranked, and probability distributions are shown as grey shaded ridges. 88 Braak-associated CpGs identified as significantly associated in bulk PFC by Smith et al. are treated as a predefined reference set, and absolute z-score distributions of these sites are represented by colored ridges. **d)** Differentially methylated region (DMR) analysis identifies regions previously identified as Braak-associated DMPs. Representative genome visualization plot of one such DMR at the *GPR37L1* locus. The red track shows p-value significance, and the blue track shows ES of CpGs at this DMR in the meta-analysis. The orange track shows the DMR, and the genome tracks are shown below. **e)** Leading-edge directional concordance analysis of PFC astrocytes and neurons for all Braak-associated CpGs identified at the tissue level by Smith et al. Smith-CpGs are ranked in order of decreasing absolute z-scores along the X-axis. Directional concordance between tissue-level ES and cell-type-specific ES is computed cumulatively across the ranked list and represented along the Y-axis. Concordance value of 0.5 represents random directional agreement.

Cell-type deconvolution limits the power of brain region-wide association testing because of the inherent dilution of signals from aggregated tissues to estimated cell types. In a previous meta-analysis of bulk cortical tissues (including PFC and TC), which utilized some of the same cohorts as those used in this study,^18^ Smith and colleagues generated DMP-level information in association with Braak stages. We leveraged this data to ask which cell-type contributed to the “bulk” Braak signal by performing a rank-based enrichment analysis of effect-size distributions across both cortical regions for each cell type, using region-specific brain DMPs identified by Smith et al. (*hereafter* called Smith-CpGs, or Smith-DMPs for simplicity). Relative to the overall distribution of effect sizes for each cell type, within the PFC, rank-based analysis shows that both astrocytes and neurons exhibit a significant shift towards larger effect sizes of Smith-DMPs **(Fig. 4b, Supplementary Table S11**), with a median absolute z-score shift of 0.32 and 0.47, respectively (Wilcoxon p = 6.49 x 10^-9^ and 9.10 x 10^-16^ respectively), and area under the curve (**AUC**) of 0.61 and 0.65, respectively (perm-p < 0.001 for both). Oligodendrocytes and endothelial cells do not show a significant effect-size enrichment for Smith-CpGs in the PFC. In the TC (**Fig. 4c**), the effect size distribution of Smith-CpGs was not enriched in any one cell type. The neuronal and astrocytic effect-size enrichment for Smith-CpGs within the PFC is also evident in the DMR analysis, where neurons shared at least 3 regions (Sidak p < 0.05) within genes implicated by Smith et al., including a highly significant DMR at the *GPR37L1* gene (Sidak p = 2.15 x 10^-5^) (**Fig. 4d**), and astrocytes shared one DMR at *RUNX1T1* (Sidak p = 0.019) (**Supplementary Fig. 6**). To further delineate the potential contributions of these two cell types to Braak stage, we examined how consistently the cell-type effects conformed in direction with the bulk tissue estimates from the Smith et al. study through a series of rank-based comparisons, starting with Smith-CpGs with the largest estimated effects. Leading-edge concordance analysis shows both neurons and astrocytes to exhibit high directional concordance to Smith-CpGs across the ranked distribution (**Fig. 4e**). Notably, among the strongest Braak-associated CpGs (highest ES), astrocytes showed a higher early concordance (ΔAUCC = 0.05 between astrocyte and neuronal concordance across the top 50 ranked Smith-CpGs), whereas neuronal concordance increased gradually and slightly exceeded astrocytes when the broader set of Smith-CpGs was considered.

### AD-dependent effects on age-associated DMPs vary by cell type

Epigenetic age acceleration is a known feature of AD. One recent study showed this age acceleration is driven by the overall glial population in the temporal lobe of AD patients,^44^ suggesting the existence of age-related cell-type vulnerability patterns associated with disease. Therefore, we wanted to assess whether AD-based methylation remodeling conforms, amplifies, or diverges from normal age-associated methylation programs in a cell-type-specific manner. We focused on cell- and brain region-specific age-associated CpGs (identified in our age-association analysis) and tested whether AD-associated methylation effects (ES) at these sites were concordant in direction (correlation) and magnitude (regression slope) compared to ES associated with normal aging. Strikingly, we observed region-dependent variability at age-associated sites (nominal) in AD (**Fig. 5a**). For example, while EC and TC exhibited cell-type-specific patterns, most cell types across the PFC showed low-to-moderate correlation of effect sizes between age and AD, as well as minimal ES amplification with AD.

**Figure 5.**
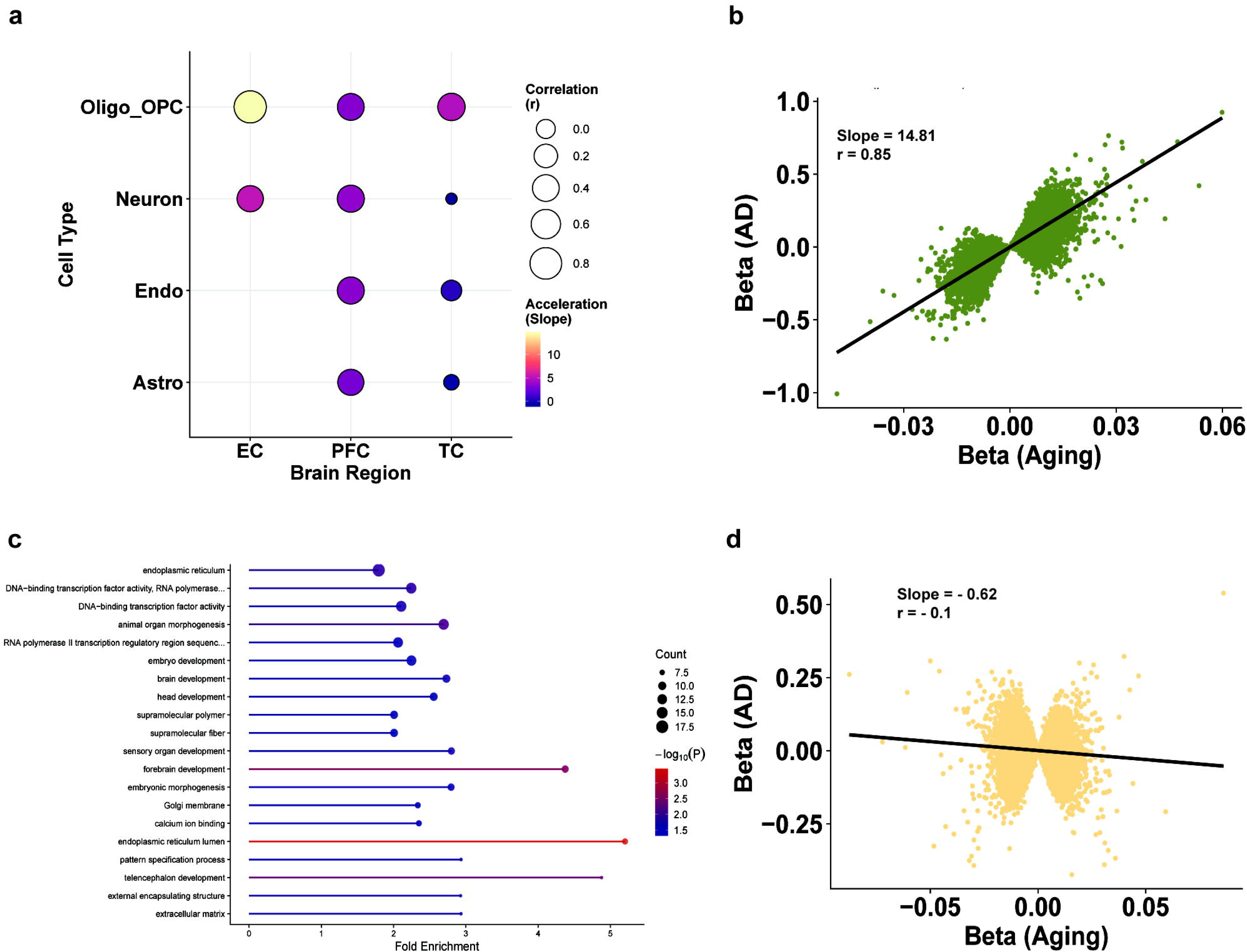
AD-dependent effects on age-associated DMPs vary by cell type. **a)** Bubble plot showing differential cell-type aging burden associated with AD status across all cell types x cortical regions analyzed. The color scale represents the slope of regression, indicative of an increase in the magnitude of ES in AD (i.e., acceleration). The size of bubbles indicates correlation of effect sizes at age-associated DMPs (p < 0.05) between age and AD associations. **b)** Correlation scatter plot of effect sizes at age-associated DMPs between age- and AD-associations in EC oligodendrocytes (r = 0.87, Slope = 0.64). **c)** Gene ontology enrichment analysis for directionally concordant age-AD CpGs at nominal significance in EC oligodendrocytes, ordered by counts. The X-axis represents fold enrichment. Color scale represents -log_10_(p). **d)** Correlation scatter plot of effect sizes at age-associated DMPs between age- and AD-associations in TC astrocytes (r = -0.05, Slope = -0.02).

For age-associated DMPs across all brain regions, the oligodendrocyte fraction exhibited moderate to high correlation (*r* = 0.4 – 0.85) between ES in age and AD, and showed an increased ES magnitude with AD. The most extreme age-to-disease ES amplification was seen in EC-oligodendrocytes (**Fig. 5b**) (slope = 14.81, r = 0.85, p < 2.2 x 10^-308^), consistent with our previous observations of shared age-AD DMPs (**Fig. 3e**), suggesting that disease-associated changes extend pre-existing age-related patterns. Of the age-associated DMPs that exhibit concordance in ES direction with AD in the EC-oligodendrocytes, gene ontology analysis shows an enrichment for broad neural development-related processes (**Fig. 5c**), suggesting an overall extension of developmental programming pathways with AD.

Interestingly, in TC, all cell types other than oligodendrocytes showed much lower amplification of ES with AD and exhibited poor correlations between age and AD ES. For example, both astrocytes and neurons show near-zero correlation between age and AD ES (*r* = -0.1 for both, p < 2.42 x 10^-46^), with astrocytes exhibiting low age-to-disease ES acceleration (**Fig. 5d**, slope = -0.62), suggesting that AD-associated methylation remodeling in these cells is not a simple extension of age-associated epigenetic programs. Tong et al. highlighted a glial signature in the temporal lobe in their cell-type epigenetic age study.^44^ We now show that this signature is likely dominated by oligodendrocytes, but not astrocytes or other cell types, revealing clear sub-glial effects in disease states.

## Discussion

In this study, we performed a comprehensive analysis of cell-type-resolved methylomic changes associated with healthy aging, Alzheimer’s disease status, and Braak stage across several brain regions. We utilized thirteen previously published AD EWAS cohorts, and integrated region-stratified meta-analysis, with rank-based effect-size enrichment and DMR discovery to uncover novel, distinct roles for glial cell types in these processes. We refine prevailing methylation models of AD and demonstrate the existence of distinct age- and disease-associated programs that fundamentally differ in scale and coherence, depending on cellular origin and anatomical context.

Across all our analyses, DNA methylation changes associated with age remained a central factor. Cell-specific effects were highly region-specific, largely dominated by glial cells within the cortex and neuronal cells in the cerebellum. Both astrocytes and oligodendrocytes showed robust age-associated methylation patterns, whereas neurons exhibited comparatively fewer sites with weaker effects in the cortex. We were able to identify 669 significant DMPs and 224 significant DMRs in PFC astrocytes that were associated with age. Although effect sizes are generally small for these individual CpG sites, larger, multi-site patterns are evident in our data and may be representative of an “aged” astrocytic phenotype. Several of these mapped to genes involved in homeostatic astrocyte function, including a highly significant DMR in the *ATP1A1* gene, involved in ion transport and synaptic communication. We also discovered several CpG sites that were differentially methylated within the *VPS52* gene, one of which was common to a previous age-EWAS of sorted brain cell types.^7^ The *VPS52* gene is a core component of the Golgi apparatus and has previously been implicated in association with *LRRK2* in Parkinson’s disease astrocytes.^45,46^ Our data now point to a novel potential role for DNA hypermethylation at sites within this gene, linking aberrant gene expression and functionality to epigenetic aging in astrocytes. To some extent, PFC astrocytes also exhibit shared aging trajectories with other brain regions, at least within the TC, consistent with the idea that glia undergo profound epigenetic remodeling with age in the cortex, reflecting changes in CNS homeostasis, metabolism, and neuronal support processes.^47,48^ Overall, our cell-type and brain region-specific analyses show distinct astrocyte and oligodendrocyte-specific methylation patterns associated with the aged cortex, and neuronal differential methylation to be associated with age in the cerebellum.

In contrast to aging, region-stratified analyses of AD case-control status revealed far fewer significant associations. Oligodendrocytes exhibit the strongest AD signal in specific cortical regions, although fewer associations were seen compared to age. Notably, we identified a novel hypermethylated DMP within the *ABCA1* gene, which is involved in cholesterol and lipid homeostasis, and is a key partner of the AD-risk factor, *APOE.*^49–51^ Dysregulation of overall *APOE* lipidation status,^52,53^ as well as accumulation of lipid droplets due to inefficient fatty acid and cholesterol metabolism in oligodendrocytes, has been previously linked with AD-relevant impaired myelin repair, white matter damage, and neuroinflammation.^54–57^ Astrocytes showed a moderate AD signal in the PFC, particularly in genes involved in calcium signaling and those encoding ion channels. Neurons, however, exhibited relatively few cortical DMPs, suggesting AD-associated differential methylation in the cortex may be driven by glia.

Despite its central role as a pathological index for AD, Braak stage produced only a handful of DMPs that survived multiple testing corrections. In analyzing region-stratified, cell-type deconvolved data, statistical power is limited. More importantly, this absence of strong Braak-associated DMPs is also likely due to the inherent nature of distributed tau pathology across the cortex, as well as cell-type resolution artifacts. In fact, Braak stage is likely highly dependent on both spatial and temporal patterning across cell types, and with modest methylation changes, these properties challenge single-CpG significance testing. To address this limitation, we employed a rank-based enrichment approach to ask whether CpG sites previously identified as significantly associated with Braak stage in bulk cortical tissues^18^ show preferential enrichment in their effect size distribution in distinct cell types. Using rank-based analysis, AUC-based enrichment, and leading-edge concordance metrics, we were able to identify clear, cell-type-specific patterns that were obscured in FDR-based single-CpG testing. Both astrocytes and neurons in the PFC showed significant effect-size enrichment of Smith-CpGs relative to their respective overall effect-size distributions across all tested CpGs, whereas no individual cell type had a pronounced effect in the TC. Astrocytes and neurons in the PFC also differed in how they were enriched for Smith-CpGs. Directional concordance analysis of these two cell types revealed a somewhat rank-dependent division of labor – astrocytes displayed a near-complete directional agreement with the top 20% of Braak-associated CpGs that marked advanced pathology (as evidenced by higher ES), whereas neuronal concordance became greater when a broader set of Braak-associated CpGs was considered. This model suggests a scenario where astrocytes align more strongly with the very highest effect Braak-CpGs, likely playing a dominant role in responding to severe pathology, exhibiting functional changes in relation to energy metabolism, homeostasis, and neuroinflammatory signaling axis.^48,58,59^ Neuronal involvement, in contrast, aligns with broader changes underlining gradual pathological remodeling across disease stages.

One key unresolved question in AD epigenomics is whether the disease-associated changes reflect a mere acceleration of normative aging programs or instead represent a distinct disease-relevant epigenetic remodeling. We find that age-associated methylation trajectories are propagated in a spatial, lineage-dependent manner. Oligodendrocytes show the strongest ES correlation of age-associated DMPs in AD, with trajectory analysis indicating an increase in magnitude of these processes across all cortical regions. Astrocytes, in contrast, show region-specific diversification. Most prominently, within the TC, age-associated ES estimates were largely not correlated with the disease state, suggesting that AD-related epigenetic remodeling in TC-astrocytes exhibits a divergence from aging-related methylomic patterns. These results support a model of lineage-defined trajectories, where oligodendrocyte aging programs define a regulatory axis along which disease progression preferentially occurs, but other cells, including astrocytes, exhibit a decoupling of age-related programs in lieu of disease-specific epigenetic remodeling.

There are some limitations to our study. To resolve cell-level DNA methylation profiles, we utilized a cell-type deconvolution paradigm that allows us to isolate cell-type signals from bulk tissue without the need for either single-cell sequencing or sorting.^24,60,61^ While this has given novel insight into cell-level dynamics in normative aging and AD-relevant disease processes, tissue-level heterogeneity, as well as disease-relevant changes in cell composition, can obscure some findings. We were also not able to assess some cell types (e.g., microglia) due to their low abundance, highlighting a need to generate more robust DNA methylation references for bulk brain tissues. Our cell-type deconvolution analysis using a region-stratified fixed-effect inverse variance weighted meta-analysis was limited in power despite having a large number of samples. Such models incur high penalties, with multiple testing corrections across several hundred CpGs that substantially reduce the effective sample size per test. This possibly limited our ability to replicate some predominantly known DMP loci associated with AD (e.g., HOXA clusters), although it is possible that these effects are dispersed across cell types, which cell-specific models cannot effectively capture. Secondly, we analyzed studies conducted on either the Illumina 450k or EPIC arrays. While these have been the gold standard for several decades, their coverage is limited, and additional studies utilizing whole genome sequencing approaches, as well as single cell sequencing methods, will be useful to uncover larger disease-relevant changes. Neuronal cells exhibit a much higher proportion of DNA hydroxymethylation compared to other cell types. However, traditional bisulfite treatment does not distinguish these bases from methylated cytosines, and as such, differential methylation in neurons reflects a potentially mixed methylation state (methylated and hydroxymethylated CpGs). This necessitates a need to further delineate methylomic “state” in these cells. Most cohorts in this study either had Braak stage data or provided only disease status. AD pathology is multivariate – future studies utilizing other pathological hallmark measures (amyloid pathology burden, amyloid-tau-neurodegeneration (ATN) scores, or Consortium to Establish a Registry for Alzheimer’s Disease (CERAD) scores) are necessary to untangle other disease-relevant associations. In this study, we showed cell-specific age-to-disease trajectories. However, since most of the obtained EWAS data were from AD-related cohorts, our age range for healthy controls was limited. Assessment of normative aging processes in the decades immediately preceding AD might be informative of whether methylomic patterns at such stages could be predictive of disease onset. While we found several relevant, novel DMP loci that implicate glial cells in aging and AD pathology, future studies should validate our findings using sorted cells, single-cell sequencing, and/or *in vitro* model systems. Lastly, we focused our study on identifying glial-specific epigenetic programs. However, endothelial cells in our analyses also exhibited some minimal epigenetic patterning, which may provide further insight into brain aging and AD pathology.

In summary, we provide a refined view of epigenetic regulatory states in AD by revealing how different cell lineages contribute to age- and disease-associated methylation patterns. We utilized a computational deconvolution method that shows brain region-specific highly abundant glial aging programs. Astrocytes within the PFC exhibit a large proportion of significant DMPs with age, whereas EC-oligodendrocytes show substantially higher DMPs in AD compared to other cell types, although sparser than those observed for age. Astrocytes and neurons emerge as key contributors to Braak-associated epigenetic remodeling in the PFC, where astrocytes show more directional concordance at loci highly associated with Braak stage and neurons exhibit a broader concordance with these disease-associated CpGs. We also uncovered a distinct age-to-disease trajectory, where oligodendrocytes exhibit a distinct age-to-disease amplification of effect sizes across all cortical regions, impacting developmental programs, but astrocytes show a divergence from age-associated epigenetic programs with AD. These insights extend tissue-level findings and underscore the importance of cell-type-resolved, rank-aware approaches for dissecting the epigenomic landscape of AD.

## Methods

### Cohorts

We identified a list of 13 archived brain-specific AD EWAS studies, consisting of a total of 3,812 independent samples (**Table 1**). Ten of these cohorts had DNA methylation data from 450k arrays, and the remaining three cohorts used EPIC arrays; all studies comprised at least 30 donors and provided publicly available data. We harmonized the probe set to assess the same CpG sites across all studies as previously described.^62,63^ The study by Brokaw and colleagues^5^ consisted of 404 unique samples for two brain regions, the middle temporal gyrus and cerebellum. Gasparoni et al. had data available for 63 and 65 individuals for the prefrontal cortex and temporal cortex, respectively.^7^ In one study by Horvath and others, data were available primarily for Huntington’s disease, but also included some individuals with AD that were included in our study; this comprised 32 samples from the cerebellum and 41 samples from the PFC.^64^ Although this study had assessed the hippocampus and temporal cortex, we did not use this data in our final meta-analysis design since it did not meet our criteria for having at least 30 samples. Lardenoije et al. had data available for the middle temporal gyrus in 82 individuals,^8^ and Laroche et al. had 242 samples from the prefrontal cortex, conducted on an EPIC array.^9^ Several of our cohorts came from archived studies in the MRC London Neurodegenerative Disease Brain Bank, including one study published by Lunnon et al. containing data for the cerebellum, entorhinal cortex, prefrontal cortex, and superior temporal gyrus,^10^ as well as one by Smith and colleagues containing cerebellum and entorhinal cortex data.^16^ We also leveraged archived cohorts in the AD knowledge portal, and assessed 546 samples in the Rush University Medical Center: Religious Order Study (ROS) and the Memory and Aging Project (MAP), i.e., ROSMAP cohort, for age- and AD-associations, as well as 734 samples for Braak stage associations in the PFC (all 450k array data).^6^ AD associations did not include samples that fell under the MCI category, but since Braak staging is a continuous measure, we included these in our latter analyses. The ROSMAP cohort was also used in a second study (RR_APOE4) to assess 254 cerebellum and prefrontal cortex samples using EPIC arrays.^65^ We also obtained EPIC array data of 223 individuals for the prefrontal cortex from the University of Pittsburgh Alzheimer’s Disease Research Center from the AD knowledge portal. Semick and colleagues had available data across four brain regions (PFC, EC, CBL, HC), each comprising at least 60 individuals that was used in this study.^14^ We also included one cohort consisting of 68 superior temporal gyrus samples,^19^ and another that contained 144 STG, and 142 PFC samples,^17^ both previously published. Since MTG and STG broadly form a part of the overall temporal cortex, we grouped these as TC to increase power for the meta-analysis. We also only included regions where we had data from at least 2 cohorts, each with a minimum of 30 individuals, and a total of at least 100 samples. In total, we were able to assess 1,626 samples in the PFC (9 cohorts), 794 samples in the TC (3 cohorts each for MTG/STG, and one for TC), 154 samples in the EC (3 cohorts), and 832 samples in the CBL (6 cohorts), each for at least 3-4 cell types.

**Table 1.**
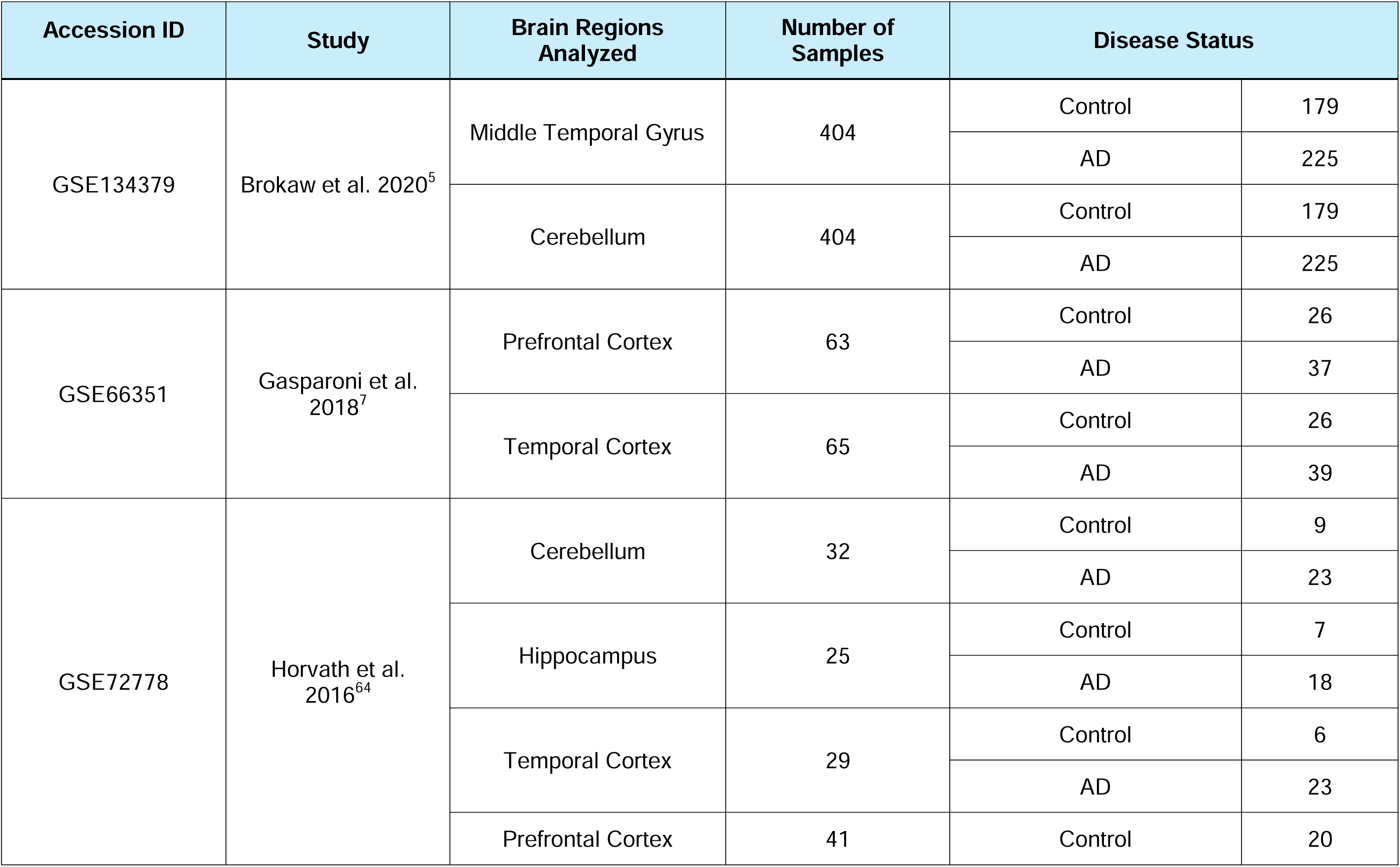
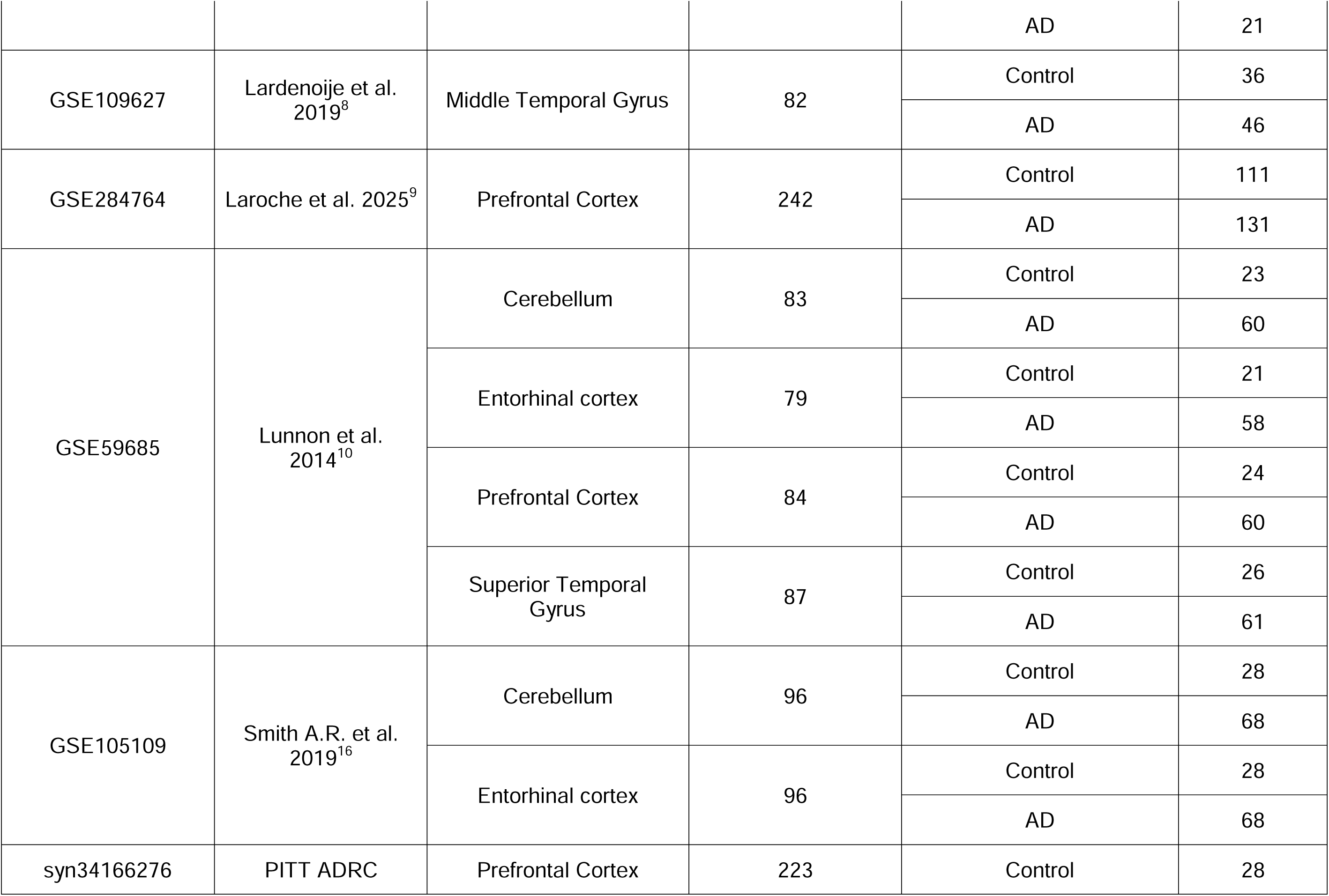
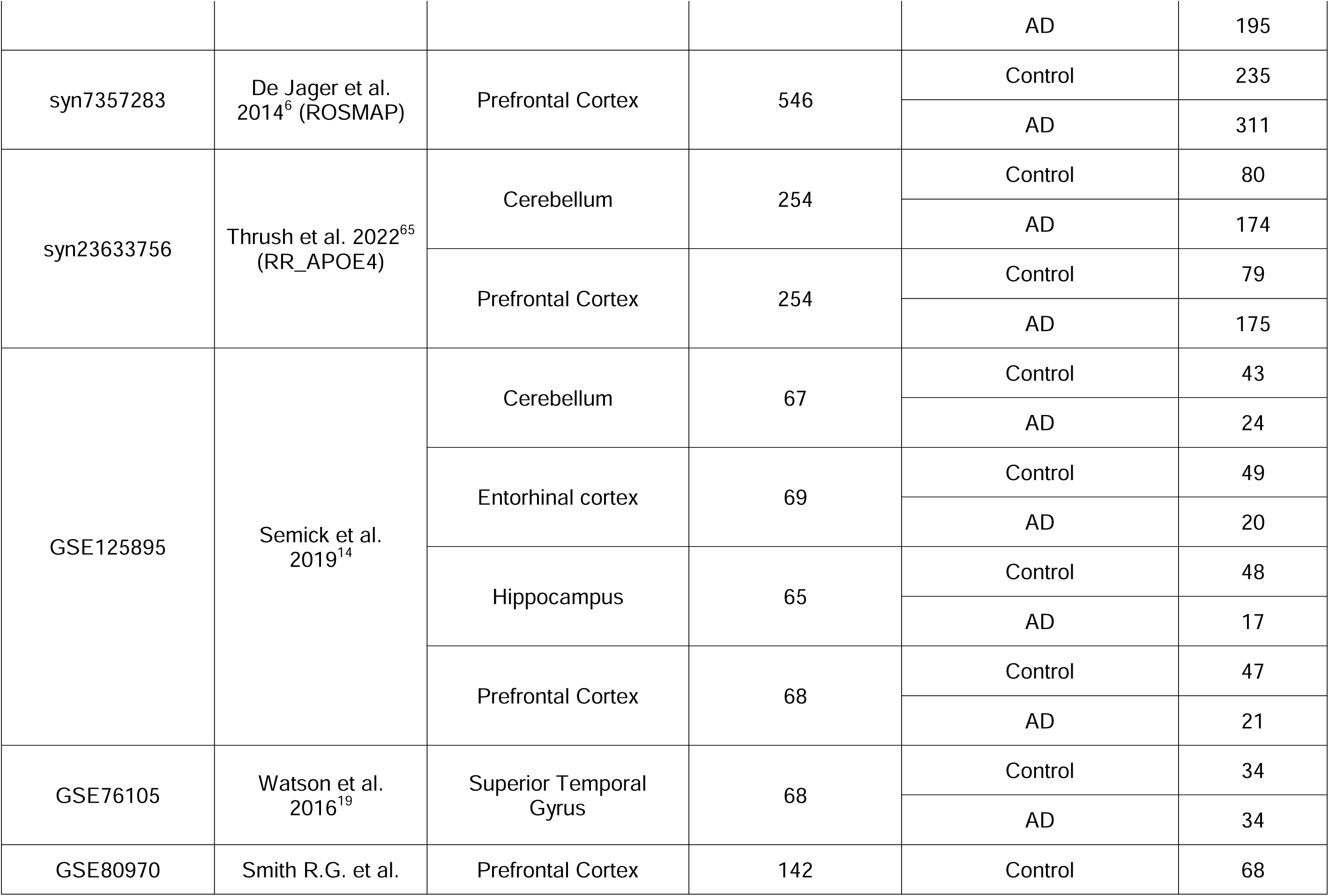
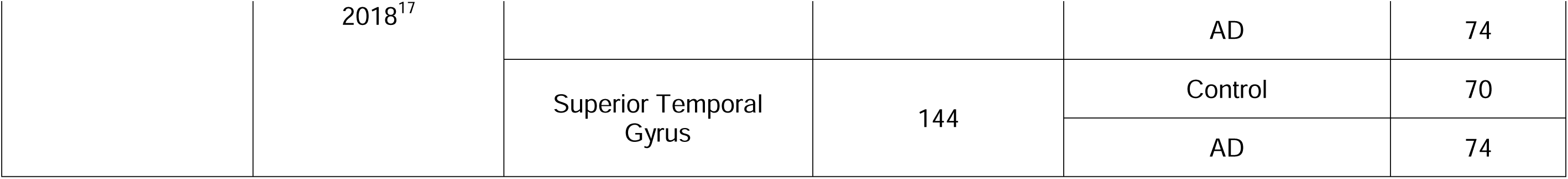
Demographic details for all cohorts used in the meta-analyses.

### Data Preprocessing

All computational analyses were carried out in R, v 4.5.1. IDAT files from raw Illumina HumanMethylation 450K or EPIC arrays for each cohort were imported and an *RGChannelSet* object was created using the *minfi* package.^66^ Sample-level quality control was performed similarly to a previous publication.^18^ Samples were removed if they had a) bisulfite conversion efficiency < 80%, b) abnormal background intensity of negative probes (< 1000), c) outlier mean methylated or unmethylated intensities (> 3 SDs), d) discordance between predicted and reported sex, or e) genetic identity mismatches detected using SNP probes where available. Probe-level QC was carried out using *minfi* and *ewastools* packages,^67^ and implemented using the ChAMP package in R.^68^ Probes present on sex chromosomes, those containing SNPs at the CpG interrogation site, and those with known cross-reactivity were removed. Background correction and dye-bias normalization were carried out using the NOOB method, implemented in *minfi*. Probes with detection *P* values > 0.01, and those with < 3 beadcounts in at least 5% of the samples were removed. Signal intensities of probes passing quality metrics were converted to beta values. In cohorts where raw IDAT files were not available, signal intensity data files were used, and a *MethylSet* object was generated using *minfi*. All other probe-level QC was performed as before, except for beadcount filtering, for which data was not available. Finally, BMIQ-based normalization^69^ was implemented on beta value matrices using the ChAMP package.

### Cell type deconvolution and differential methylation analysis

To estimate cell proportions within each cohort, we used a previously published DNA methylation reference matrix (EpiSCORE) for brain tissues,^26^ that was implemented in a prior deconvolution study.^24^ Bulk data was processed for each brain region separately. First, the constAvBetaTSS function from the EpiSCORE package was used to map CpG loci to Entrez IDs to derive promoter-level methylation estimates. Cell proportions were then estimated using a weighted robust partial correlation method, implemented in EpiSCORE. The reference matrix in EpiSCORE can be used to estimate proportions for at least six unique brain cell types. Because we detected < 0.5% of microglia across several cohorts, these were dropped from further analysis. Oligodendrocyte estimates were relatively small, and therefore, we grouped these with OPCs as done previously to allow better power for the detection of significant associations.^24^

Cell-type-specific association analysis for age, AD, and Braak stage was performed using the CellDMC framework, implemented in the EPIDISH package.^27,70^ This method models interactions between estimated cell type proportions and phenotypic variables to identify CpGs exhibiting differential methylation effects attributable to a given cell type. To do this, we first harmonized CpG probes to those existing on both the 450k and EPIC arrays, then prespecified association models for testing. Briefly, for age, AD status, and Braak stage, we performed region-stratified analysis with criteria for a minimum sample size of 30 per cohort, with > 5 per outcome group and > 5 per region. For age associations, we controlled for sex and diagnosis as covariates and modeled age as a continuous variable. For AD, we used age and sex as covariates, modeling AD diagnosis as a binary variable and Braak stage as a continuous variable. To control for unmeasured technical and biological confounders, we used surrogate variables (**SV**), implemented using the *sva* package.^71^ SVA models included the outcome of interest (age, AD, or Braak stage), covariates, and cell-type fractions. The number of SVs was selected adaptively by iteratively increasing the number of SVs until the genomic inflation factor (λ) for the given outcome term per region was reduced to ≤ 1.2, as previously implemented by Smith et al. ^18^ If baseline inflation was already below the threshold, no SVs were included for that model. Finally, CellDMC models were fit using beta values as the response variable, with phenotype terms, covariates, and selected SVs included as predictors. Cell-type specific effect sizes, standard errors, and p-values were extracted for each CpG, and multiple testing correction was applied using both the Benjamini-Hochberg false-discovery rate method,^72^ and Bonferroni correction.^73^ We mainly reported FDR-based significance metrics in this study, although Bonferroni adjustments are available upon request. Unless otherwise specified, FDR < 0.05 was used as a statistical significance threshold for all analyses. For each model and cell type, genomic inflation was computed after association tests, and quantile-quantile plots were generated to assess residual test statistic inflation, and association results eligible for meta-analysis were exported for further processing.

### Region-stratified cell-specific meta-analysis

Cell-type-specific association results from individual cohorts were meta-analyzed separately for age, AD status, and Braak stage, using an inverse variance fixed-effects framework. Meta-analysis was performed at the level of CpG x region x cell type, by integrating ES and SEs derived from the CellDMC output. CpG-level results were required to be supported by at least two independent cohorts to be eligible for meta-analysis. To control for residual test statistic inflation and bias at the cohort-region-cell type level, effect sizes were corrected using the BACON method before meta-analysis.^74^ Groups of phenotype x region x cell type were excluded from further analysis if genomic inflation exceeded 2.0 before BACON correction, or remained > 1.5 after correction, ensuring that only well-calibrated cohorts contributed to the meta-analysis. We implemented the inverse-variance weighted fixed-effects model via the *metagen* function within the Meta package.^75^ Random-effects models were not included for primary inference due to the typically small number of contributing cohorts; however, heterogeneity statistics (Cochran’s Q, I^2^) are reported. FDR significance was set at 0.05.

### Differentially Methylated Region (DMR) Analysis

Differentially methylated regions were identified from cell-type-specific meta-analyses results using the *comb-p* method^76^ implemented in Python (v 2.7) as previously described. ^18^ Regions were spatially aggregated within a 500bp window and seeded using a nominal CpG p-value threshold of 0.05, then corrected for multiple testing using the Sidak method. DMRs were annotated to nearby genes and genomic features (*hg19*), and representative loci were visualized using genomic context tracks, implemented using the *Gviz* package in R.^77^

### Effect-size rank enrichment analysis

To evaluate cell-type contributors to tissue-level Braak stage DMPs previously reported by Smith et. al.,^18^ we used a rank-based enrichment analysis of effect-sizes within each region and cell type. For each cell-type per region, all ∼400,000 CpG sites were ranked by their absolute z-score (defined as ES/SE)_of the Braak association from the meta-analysis. Smith-CpGs were treated as a predefined reference set, and enrichment was assessed by comparing the distribution of their absolute z-scores to that of all other CpGs using the Wilcoxon rank-sum tests and permutation-based AUC statistics (1 x 10^4^ permutations). This approach assesses whether previously reported tissue-level Braak-associated CpGs tend to exhibit a stronger cell-type-specific Braak effect than expected by chance. Results were visualized using ridge density plots.

To further characterize the directional concordance between Smith-CpGs and cell-type-specific Braak associations (PFC astrocytes and neurons), we performed leading-edge concordance analysis. Smith-CpGs were first ranked in order of decreasing absolute z-scores, such that the CpGs with the strongest reported Braak-associations were evaluated first. Directional concordance was then computed cumulatively across the ranked list. At each rank threshold *k,* concordance was defined as the proportion of top *k* CpGs whose ES had the same direction in the cell-type-specific meta-analysis as previously reported by Smith et. al. Concordance values therefore range from 0 to 1, with 0.5 representing random directional agreement. Cell-type differences within the leading edge were evaluated using permutation-based AUCC tests, comparing directional concordance over the top 50 CpGs (1,000 permutations). Differences in directional concordance between the two cell types were interpreted qualitatively as reflecting differences in how strongly or broadly tissue-level Braak-associated CpGs align directionally with cell-specific Braak effects across the ranked sites.

### Cell-specific age-to-disease trajectory analysis

Age-to-disease trajectories were assessed by comparing direction and effect sizes of CpGs nominally significantly associated with age (p < 0.05) between cell-type-specific age and AD associations. Effect size amplification was quantified as the slope of the regression between age and AD ES estimates for the age-associated DMPs. Pearson correlation was also computed to test the extent to which the same CpGs contributed to both age and disease associations in a given cell type and region. Concordance indicates that the AD effects were in the same direction as the age effects, while ES scaling (regression) indicates whether AD-associated changes are comparable to or exceed age-associated changes.

### Gene ontology, pathway, and other visualizations

CpGs were annotated using the Illumina (UCSC) gene annotation database using the *minfi getAnnotation* function. Annotations are based on their proximity to known transcription start sites in the genome. Functional enrichment of CpGs identified in the meta-analysis was performed using the *gometh* function, implemented in the *missMethyl* package in R.^78^ All other data visualizations, including summary plots, Manhattan plots, and volcano plots, were generated using the *ggplot2* package in R.

## Acknowledgements

This work was supported through the National Institute of Aging of the National Institutes of Health, under award number R21AG085428 (to M.A.C.). The authors acknowledge the computational support received from the High-Performance Computing Center (ARC) at the University of Texas at San Antonio (UTSA) and data storage support from UTSA’s hyper-converged infrastructure, operated by University Technology Solutions. We thank the authors of all prior publications that made their data publicly available, making this analysis possible, as well as the donors and families who made this research possible.

## Author Contributions

U.B. undertook data analysis and bioinformatics. U.B., M.Z.K., and M.A.C. conceived the idea and developed the project. M.A.C. provided funding support. U.B., M.Z.K., and M.A.C. drafted the manuscript. All authors read and approved the final submission.

