## Supplementary figures and images for "A Cell-Type–Resolved Meta-Analysis Reveals Glial DNA Methylation Changes Associated with Aging and Alzheimer’s Disease"

### Figure S1

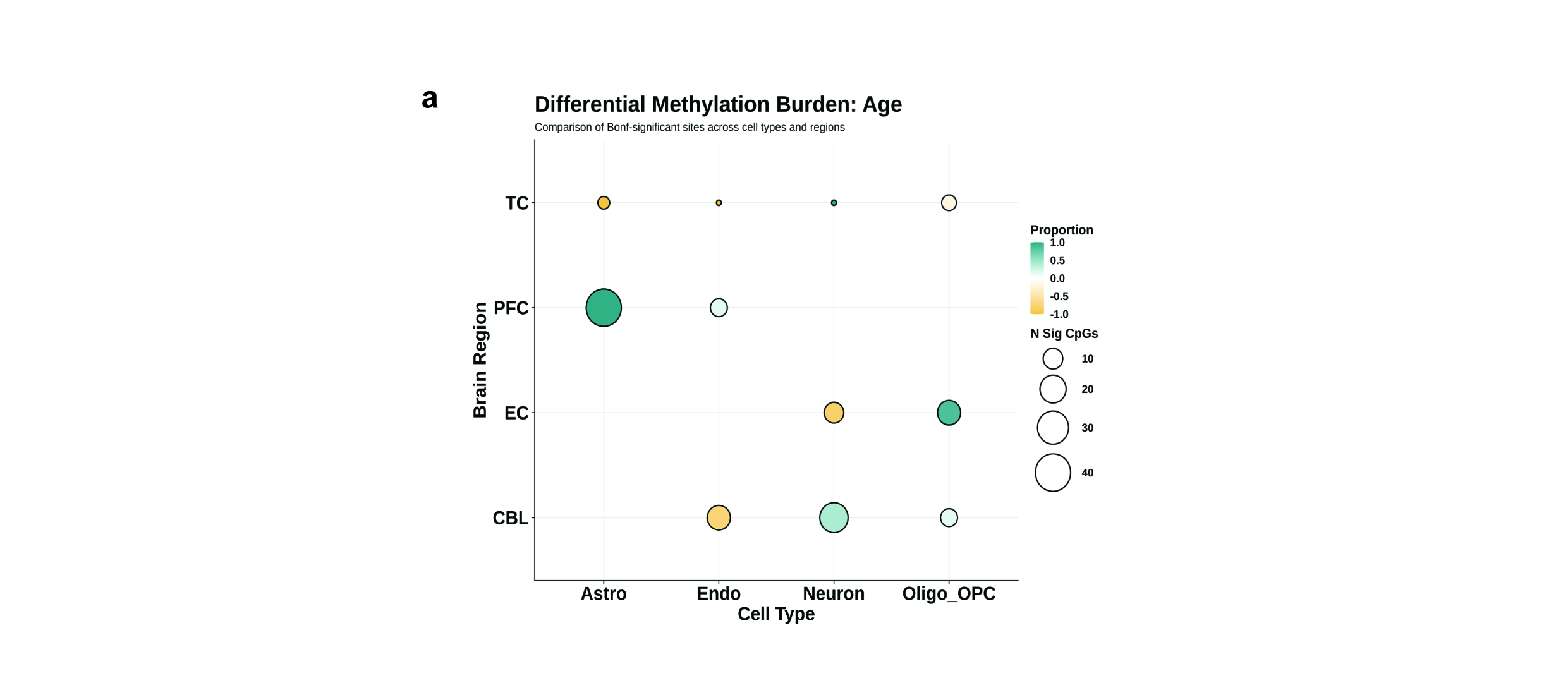

### Figure S2

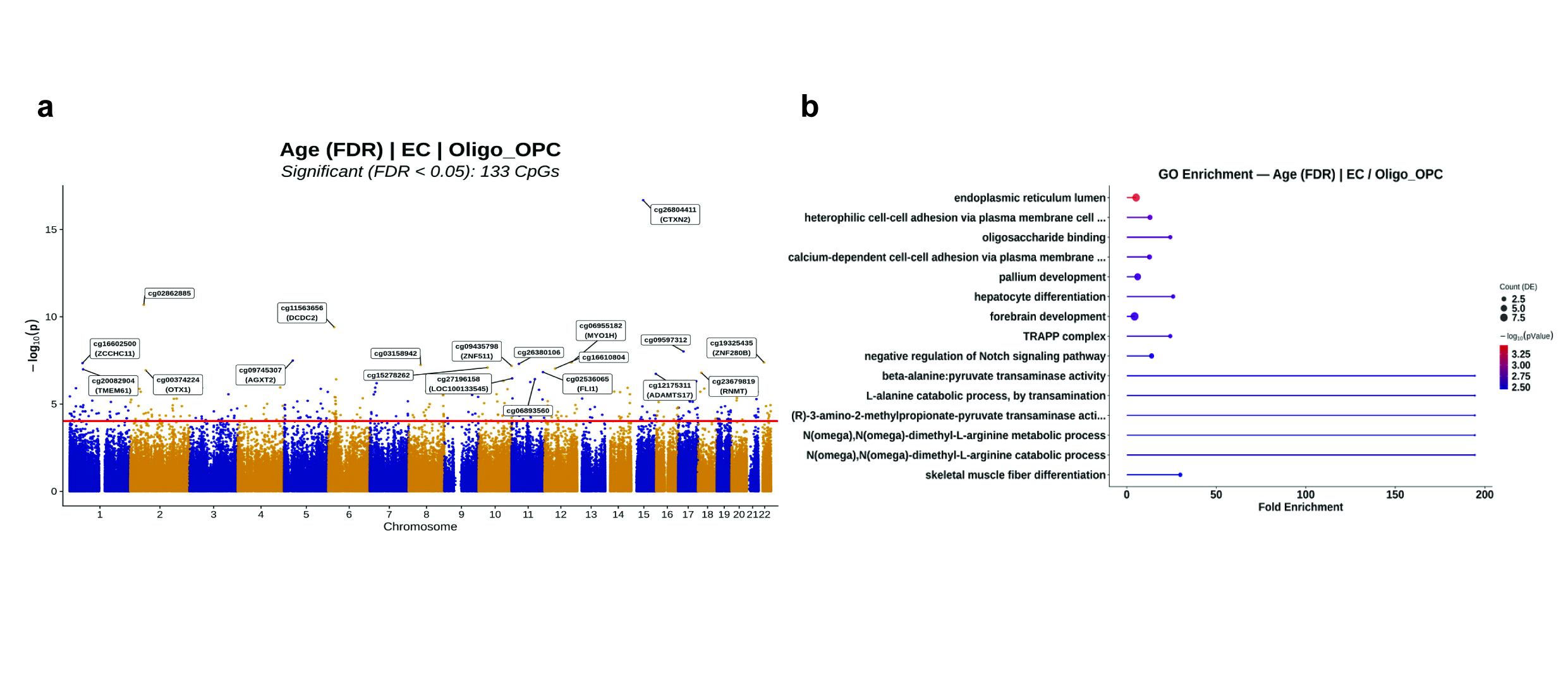

### Figure S3

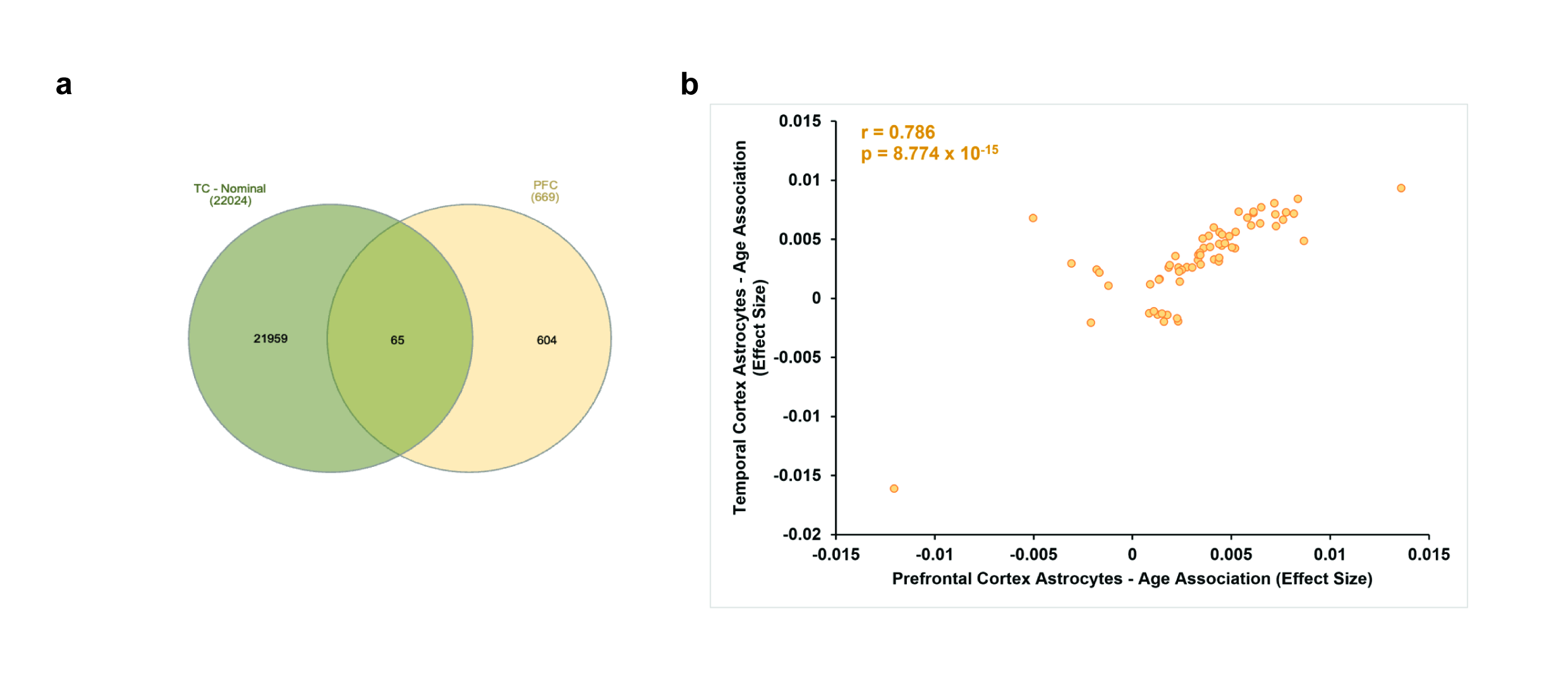

### Figure S4

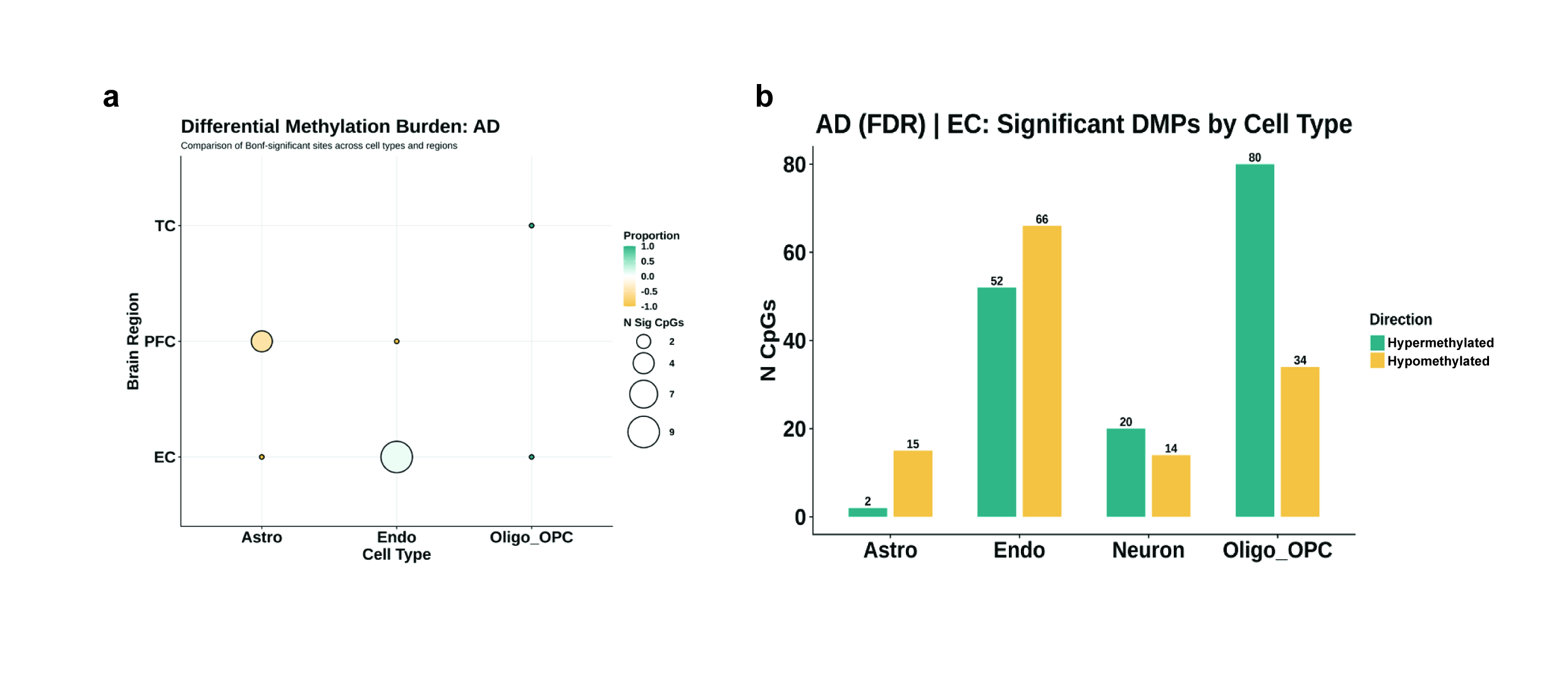

### Figure S5

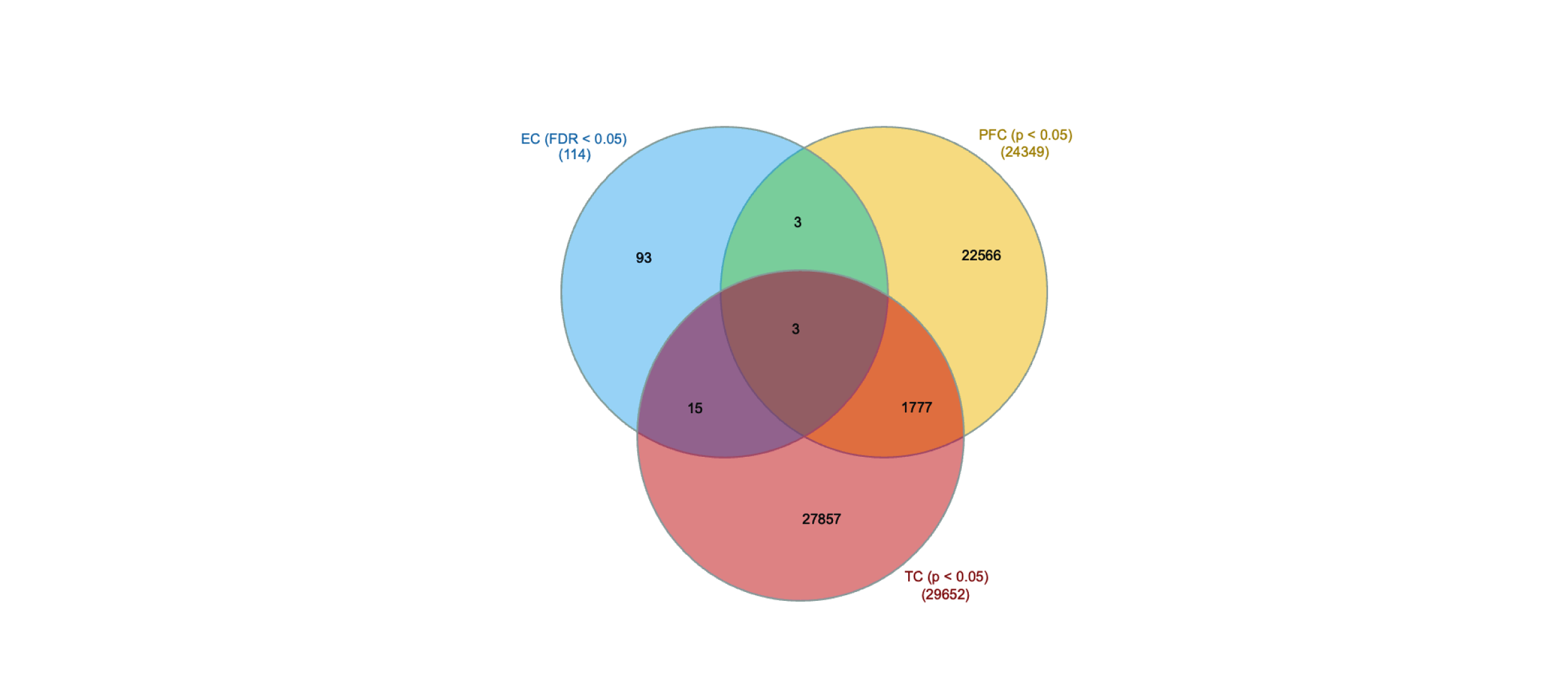

### Figure S6

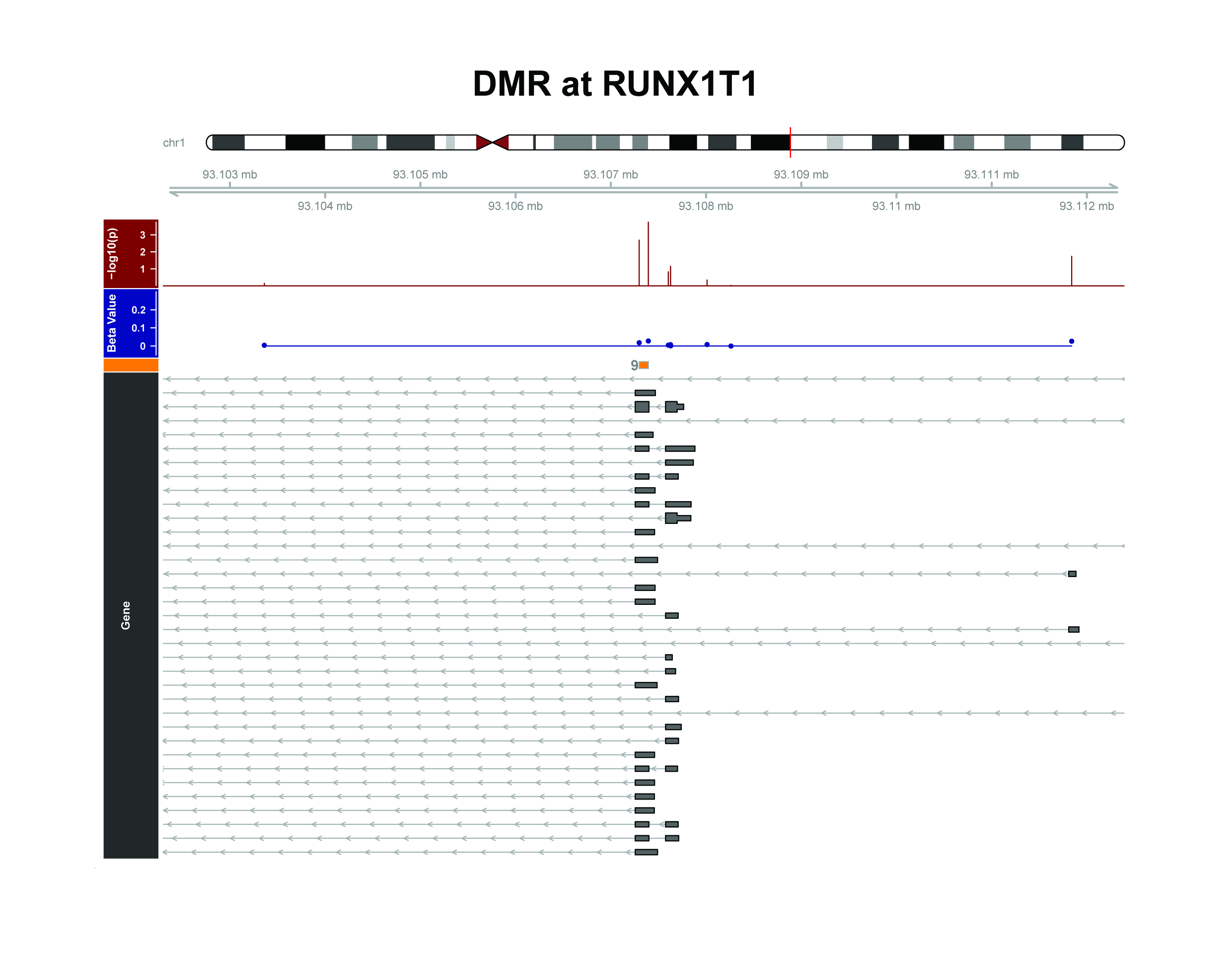
